# Holobiont Evolution: Mathematical Model with Vertical *vs.* Horizontal Microbiome Transmission

**DOI:** 10.1101/465310

**Authors:** Joan Roughgarden

## Abstract

A holobiont is a composite organism consisting of a host together with its microbiome, such as a coral with its zooxanthellae. Hosts and their microbiomes are often intimately integrated in behavior, physiology, anatomy, and development. To explain this integration, some investigators contend that selection operates on holobionts as a unit and view the microbiome’s genes as extending the host’s nuclear genome to jointly comprise a hologenome. Holobiont selection then operates on holobionts with their hologenomes by analogy to how ordinary natural selection operates on individuals with their genes. Other investigators point out that vertical transmission of microbiomes is uncommon. They contend that holobiont selection cannot be effective because a holobiont’s microbiome is an acquired condition rather than an inherited trait. The disagreement between these positions invites a simple mathematical model to see how holobiont selection might operate and to assess its plausibility as an evolutionary force.

Here I present two variants of such a model. In one variant, juvenile hosts obtain microbiomes from their parents (vertical transmission). In the other variant, microbiomes of juvenile hosts are assembled from source pools containing the combined microbiomes of all parents (horizontal transmission). According to both variants, holobiont selection indeed causes evolutionary change in holobiont traits. Therefore, holobiont selection is plausibly an effective evolutionary force with either mode of microbiome transmission.

Furthermore, the modeling employs two distinct concepts of inheritance, depending on the mode of microbiome transmission: collective inheritance whereby juveniles inherit a sample of the collected genomes from all parents as contrasted with lineal inheritance whereby juveniles inherit the genomes from only their own parents. Collective inheritance may also apply to the evolution of soil and other microbes that feature large amounts of horizontal gene transfer and may underlie cultural evolution wherein each generation inherits a sample from the collected knowledge of the preceding generation. A distinction between collective and lineal inheritance also features in theories of multilevel selection.

## Introduction

This paper develops a simple conceptual mathematical model for the evolution of holobionts. Its purpose is to clarify how holobiont selection may cause evolutionary change in the traits of holobionts.

A *microbiome* is an “ecological community of commensal, symbiotic, and pathogenic microorganisms” that shares the “body space” of a host (Lederberg & McCray 2001). A *holobiont* is a composite organism consisting of a host together with its microbiome (Margulis 1991). The *hologenome* is the union of all the host genes with all the genes in its microbiome (Zilber-Rosenberg and Rosenberg 2008). To these definitions I add the following: a *hologenotype* is the configuration of the hologenome in an individual holobiont. The *hologene pool* is the set of all hologenotypes in a population of holobionts. A *holophenotype* is a hologenotype’s expression in the behavior, physiology, and morphology of a holobiont. *Holobiont selection* is the differential reproduction and/or survival of holobionts as a function of their holophenotypes. The hologene pool is represented by the total holobiont population size together with a frequency distribution of hologenotypes over the space of all possible hologenotypes. Through time, holobiont selection changes both the total number of holobionts as well as the frequency distribution of hologenotypes.

Many researchers have suggested that selection operates on the combination of the host together with its microbiome as a unit (Woese 2004, Rosenberg, Koren, Reshef, Efrony, & Zilber-Rosenberg 2007, Fraune & Bosch 2010, Guerrero, Margulis & Berlanga 2013, Gilbert, Sapp & Tauber 2012, McFall-Ngai, Hadfield, Bosch, Carey, Domazet-Loo, Douglas, Dubilier et al. 2013, Rosenberg & Zilber-Rosenberg 2014, Bordenstein & Theis 2015, Lloyd 2017, Rosenberg & Zilber-Rosenberg 2018). According to this perspective, the union of a host’s genes together with the genes from all the microbes living in it comprises an extended genotype, called here the hologenotype. Selection on holobionts then changes hologenotype frequencies in a population of holobionts with a process envisioned as similar to how classic natural selection on individuals changes genotype frequencies in a population of individuals. Proponents of holobiont selection have documented astonishingly intricate and intimate integration between hosts and their microbiomes in behavior, physiology, anatomy, and development and argue that this integration itself is evidence of the power of holobiont selection to produce host/microbiome coadaptation.

Other researchers are unconvinced that holobiont selection actually accomplishes the evolutionary results claimed for it (Booth 2014, Moran & Sloan 2015, Douglas & Werren 2016, Hester, Barott, Nulton, Vermeij, & Rohwer 2015, Skillings 2016, Chiu & Eberl 2016, Queller & Strassmann 2016, Doolittle & Booth 2017). They observe that vertical transmission of microbiomes from parental hosts to juvenile hosts is uncommon. Instead, juvenile hosts usually acquire microbiomes from the surrounding environment leading to horizontal transmission. Hence, the microbiome in a holobiont is an acquired trait. Because the microbiome is not inherited, holobiont selection presumably cannot cause holobiont evolution.

The skeptics acknowledge the existence of host/microbiome integration, but deny that holobiont selection accounts for the evolution of this integration. They note further that microbiomes are not necessarily helpful to the host, but can contain pathogens that hurt the host. They argue that a balanced view of the host/microbiome relation would include more mention of deleterious microbes to supplement the mutualistic microbes emphasized by the holobiont selection proponents.

The response of holobiont selection proponents to the objections raised by the skeptics is to demonstrate that vertical transmission is actually more common that the skeptics claim. The microbiome does not need to be contained in the gametes of the parental host to be considered as being transmitted vertically. The parental host’s microbiome may colonize (or infect) the juveniles during birth, for example during passage through the birth canal in mammals, or from contact with nest material in birds, and so forth.

Although the skeptics may underestimate the amount of vertical transmission of microbiomes, this response by the proponents still remains unsatisfying. For example, in a few species of corals the zooxanthellae are transmitted vertically in the coral’s gametes (Hirose & Hidaka 2006). But the vast majority of zooxanthellae are acquired from the open sea water surrounding the coral (Babcock, Bull, Harrison et al. 1986, Trench 1993). Similarly, vertical transmission of the microbiome in sponges is described as inconsistent and unfaithful (Björk J, Díez-Vives C, Astudillo-Garćıa C, Archie1 EA, & Montoya J. 2019). Although both these examples are marine, even terrestrial groups may offer less vertical transmission than might be expected. In a live-bearing cockroach, researchers found only one component of the microbiome to be transmitted vertically, a bacterium that apparently supplies nutrients needed during development (Jennings 2019), but also found that a majority of the microbiome components are acquired from the environment throughout the first-instar and melanization period (Jennings EC, Korthauer MW, Hamilton TL & Benoit JB. 2019). So, the limited vertical transmission of the microbiome in many types of holobionts remains a difficult point for holobiont selection proponents.

On the other hand, the holobiont selection skeptics face the reality that extensive and intri-cate host/microbiome integration has indeed evolved somehow. The skeptics argue that classic coevolutionary selection operating on separate, yet interacting, host and microbiome genotypes can produce that integration. It is well known that coevolutionary selection has led to many co-adaptations between macro-organisms such as the classic plant-pollinator and damselfishsea anemone associations, among others. Coevolutionary selection has also produced intricate schemes by which hosts reduce parasite virulence involving both chemical mimicry and geo-graphically located arms races (Nash, Als, Maile, Jones, & Boomsma 2008). Furthermore, both horizontal and vertical pathogen transmission leading to the coevolution of reduced virulence by hosts has been studied theoretically (Yamamura 1993, Lipsitch, Siller, & Nowak 1996). Still, the degree of integration between host and microbiome is claimed by holobiont proponents to be of an entirely different scale than that seen in the known examples of coevolutionary coadaptation. This opinion, however, is subjective and skeptics can assert that given enough time, classic coevolutionary selection will eventually yield the degree of host/microbiome integration highlighted by the proponents.

So, at this time, two quite reasonable alternative positions occur with respect to holobiont selection. The proponents face the difficulty of dealing with the largely horizontal mode of microbiome transmission. The skeptics must insist that classic coevolutionary selection is sufficient to produce the deeply integrated coadaptation that occurs in the host/microbiome relation.

In this context, a simple conceptual model might help clarify how holobiont evolution can be attained with holobiont selection. This article specifically addresses the question of microbiome transmission. It analyzes whether holobiont evolution *via* holobiont selection is plausible depending on the mode of microbiome transmission. This study responds to the demand from medical and conservation constituencies for understanding the microbiome/host relationship (Turnbaugh, Ley, Hamady, Fraser-Liggett, Knight & Gordon 2007, Dubilier, McFall-Ngai & Zhao 2015, OSTP 2016).

The models introduced here are intentionally simplistic. Many review articles document an extraordinary complexity of microbiomes across diverse systems in mammals, plants, soils, and oceans of interest to medicine, agriculture and conservation. The modeling task is to distill this detail into a minimum set of assumptions relevant to conceptual clarity, not to empirical completeness. For example, two versions of model are introduced. In one, almost all of the microbiome is transmitted vertically with but a trickle of horizontal mixing; in the other all the microbiome is transmitted horizontally without any vertical transmission. In reality of course, both modes occur simultaneously. That empirical truth is not relevant here, however. If holobiont selection is effective with near total vertical transmission and also with total horizontal transmission, then there is little reason to doubt it will also be effective if both modes of transmission occur simultaneously. It is sufficient for conceptual clarity to show that holobiont selection works in the two extreme cases, and to leave formulation of a model where some fraction of the transmission is vertical and the remaining fraction horizontal to the future should the need for such an elaborate model arise. I hope the reader will appreciate that simple models such as these are always vulnerable to attack for leaving out some cherished detail, and will grant me license to concentrate on assumptions that seem most relevant to bringing conceptual clarity to the holobiont *v*s. coevolutionary selection dispute. The models in this article are minimum models for concepts, not minimum models for empirical systems.

## The Hologenome

Because the hologenome consists of the genes from all the microbes in the microbiome plus the host’s nuclear genes, the mere description of the hologenome might seem daunting. The approach introduced here is to represent a hologenome as a hierarchy.

Figure 1 diagrams two instances of a conceptual holobiont, *A* and *B*. A representative host is depicted as a rectangle with rounded corners. A host’s genome contains nuclear genes depicted in the gray gear box. Holobiont-*A* has allele *H_1_* and Holobiont-*B* has *H_2_* at a locus of interest in its host’s nucleus. Each holobiont has two niches for microbes depicted as green or brown that can represent different physical locations within the host or different functional roles. Two taxa of greens, circle and star, and two taxa of browns, rectangle and diamond, are available for each niche. The abundance of these taxa in each niche can vary among holobionts—Holobiont-*A* has three circles and Holobiont-*B* two stars. The allelic composition can vary among microbes—the circles have one *C_1_* allele and two *C_2_* alleles in Holobiont-*A*.

**Figure 1:**
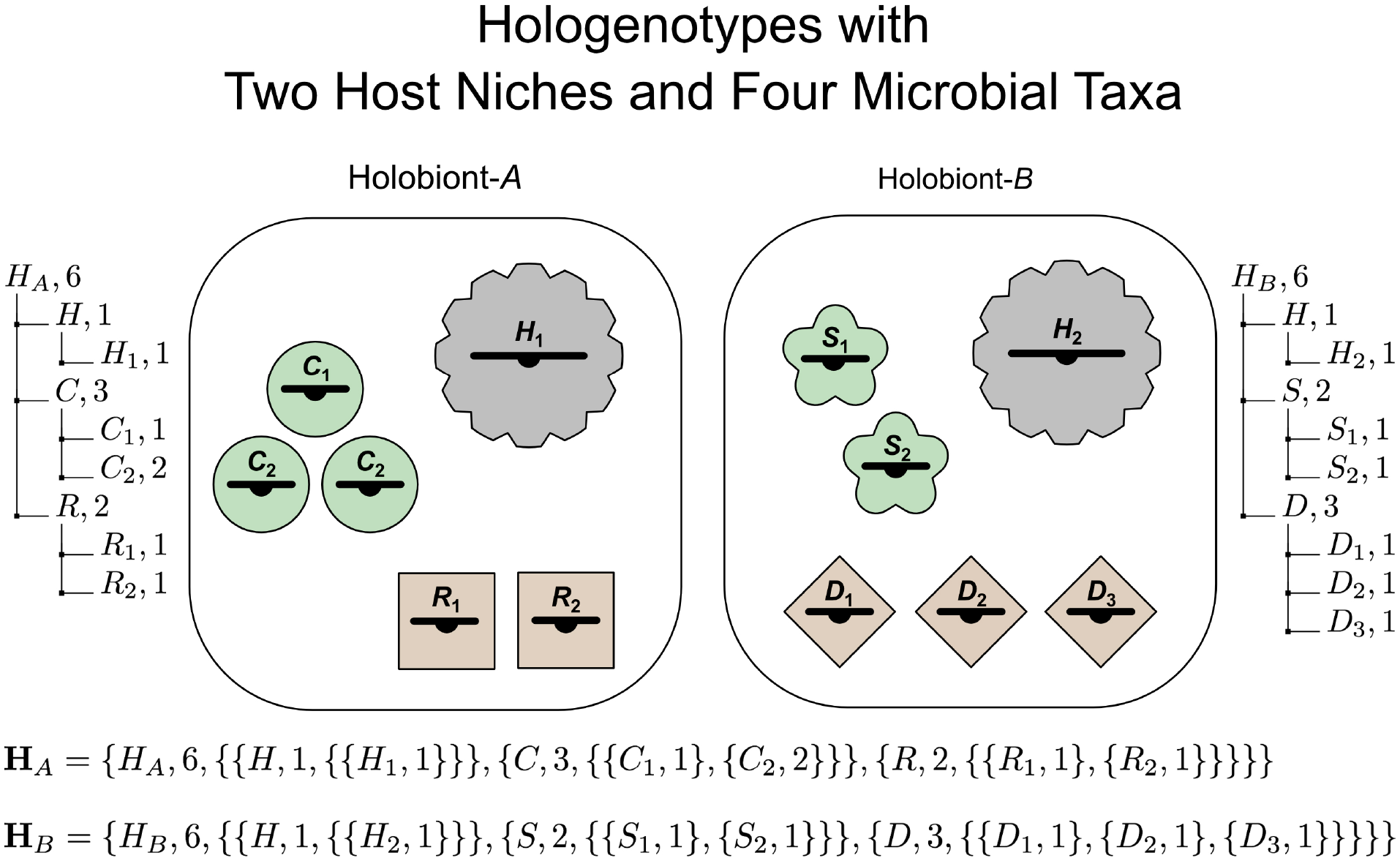
Examples of hologenotypes from two hosts assuming two microbe niches (green and brown) and two possible microbe taxa per niche (circles and stars, rectangles and diamonds) together with allelic genetic variation within the microbe taxa. The hologenotype may be represented as a list of lists, as illustrated at bottom of figure. The right and left sides of the figure illustrate the genetic hierarchy for the hologenotypes. **H**_A_, for example, reads as consisting of six individual genomes, one genome being the host nucleus containing one H_1_ allele, three genomes from circle microbes containing among them one C_1_ allele and two C_2_ alleles, and two genomes from rectangle microbes containing among them one R_1_ allele and one R_2_ allele.

The hologenotype of an individual holobiont may be represented as a hierarchical data structure (list of lists) as depicted in Figure 1. The outer list indicates the number of genomes in the holobiont. (The host nucleus is counted as a taxon containing one genome.) Nested within this are lists for the abundance and identity of each genome. And nested within this are lists for the number and identity of each allele.

In the hologenotype description the number of microbes is normalized to a standard host cell. For example, if Holobiont-*A* as a whole is comprised of 1000 host cells, 3000 green circle microbes and 2000 brown rectangle microbes, then its hologenotype description, **H***_A_*, records 1 host nucleus, 3 green circles, and 2 brown rectangles as shown in Figure 1.

A model of holobiont evolution predicts the trajectory through time of the hologenotype numbers for a holobiont population.

Burke, Steinberg, Rusch, Kjellberg & Thomas (2011) have suggested that a bacterial community should be described in terms of bacterial functions rather than bacterial species. The hologenotype description in Figure 1 includes this possibility with its use of niches. A niche can be a region of the host, say gut *vs.* mouth *etc.* But a niche can also be a functional role that can be played by any of several bacterial species. This later interpretation accords with common usage in ecology where community structure is also described in terms of niches—in island lizards in the Caribbean for example, distinct functional niches refer to species who feed in grass fields and on bushes, on tree trunks near the ground, higher on tree trunks, in the central canopy of trees and on canopy twigs. These characteristic niches are populated by different species on different islands and in different habitats yielding a recurring community structure from place to place even though the species identities in those places vary (*cf.* Roughgarden 1995, Losos 2011). The suggestion that bacterial communities should be described in terms of bacterial functions rather than species accords with how community ecology has long described community structure in terms of niches.

Similarly, Doolittle and Booth (2015) and Doolittle and Inkpen (2018) have argued that selection pertaining to microbiomes should focus on the biochemical functions of bacteria rather than on their specific taxa. They write that biochemical functions may be conserved across hosts and through time whereas the taxa preforming those functions are variable. They argue that the functions themselves rather than the taxa should be considered as the objects of selection. This perspective is also accommodated with the type of hologenotype description illustrated in Figure 1. A hologenotype with say, no green taxa could be matched against a hologenotype with no brown taxa. Holobiont selection in a holobiont population with these two hologenotypes would yield a result about the evolution of the niches themselves. Moreover, multiple functionally equivalent taxa could populate a niche, facilitating an investigation of within-niche species diversity based on within-niche neutral species.

Gene duplication is also covered by the hologenotype description of Figure 1. One can represent a duplicated nuclear gene with say, two copies of the *H_1_* allele in the the gray gear box of one hologenotype matched against one copy of the *H_1_* allele in the gray gear box of another hologenotype. Holobiont selection in a population with these two hologenotypes would reveal the evolutionary outcome of nuclear gene duplication.

Moreover, microbe abundance is in fact the equivalent of gene duplication—duplication of microbe genes rather than host nuclear genes. A hologenotype with say, two *C_1_* circle microbes could be matched against another hologenotype with say, three *C_1_* circle microbes. Holobiont selection with these two hologenotypes amounts to selection for the microbe’s allele copy number. Alternatively, this selection could be viewed as selection for microbe abundance. Indeed, the role of the microbes can be looked at from either of two sides, as though an Escher painting. From the hologenome standpoint, the microbes are no more than encapsulated genes whose abundance represents a gene copy number. From a microbiology standpoint, the microbes are tiny living entities whose abundance represents the community structure of a microbiome. From the hologenome standpoint, holobiont selection is affecting gene copy number whereas from the microbiology standpoint, holobiont selection is affecting the microbiome community structure.

Thus, the hologenotype description of Figure 1 is very general and supports the future modeling of many evolutionary scenarios should the need arise. The goal in this paper is to compare the evolutionary outcomes of vertical and horizontal transmission. For this purpose it is sufficient here to analyze the special case of hologenotypic variation solely in microbe number, assuming one host niche, one microbial taxon, and no allelic variation in host or microbe genomes. The microbe may be either a pathogen or a mutualist.

First, the case of vertical transmission is developed, followed by the case of horizontal transmission.

## Vertical transmission

Three processes occur as sequential stages within each host generation, as shown in Figures 2 and 3 (Roughgarden 2017, Roughgarden, Gilbert, Rosenberg, Zilber-Rosenberg and Lloyd 2018). The host’s generation time is the macro time step. The macro time step begins in the upper right of the figures, *i.e.* right after the holobiont selection stage and before the mixing stage. Within each macro time step, micro time steps reflect the kinetics of microbe exchange and microbe generation time. The text offers a verbal description of the model while the equations are found in the mathematical appendix.

**Figure 2:**
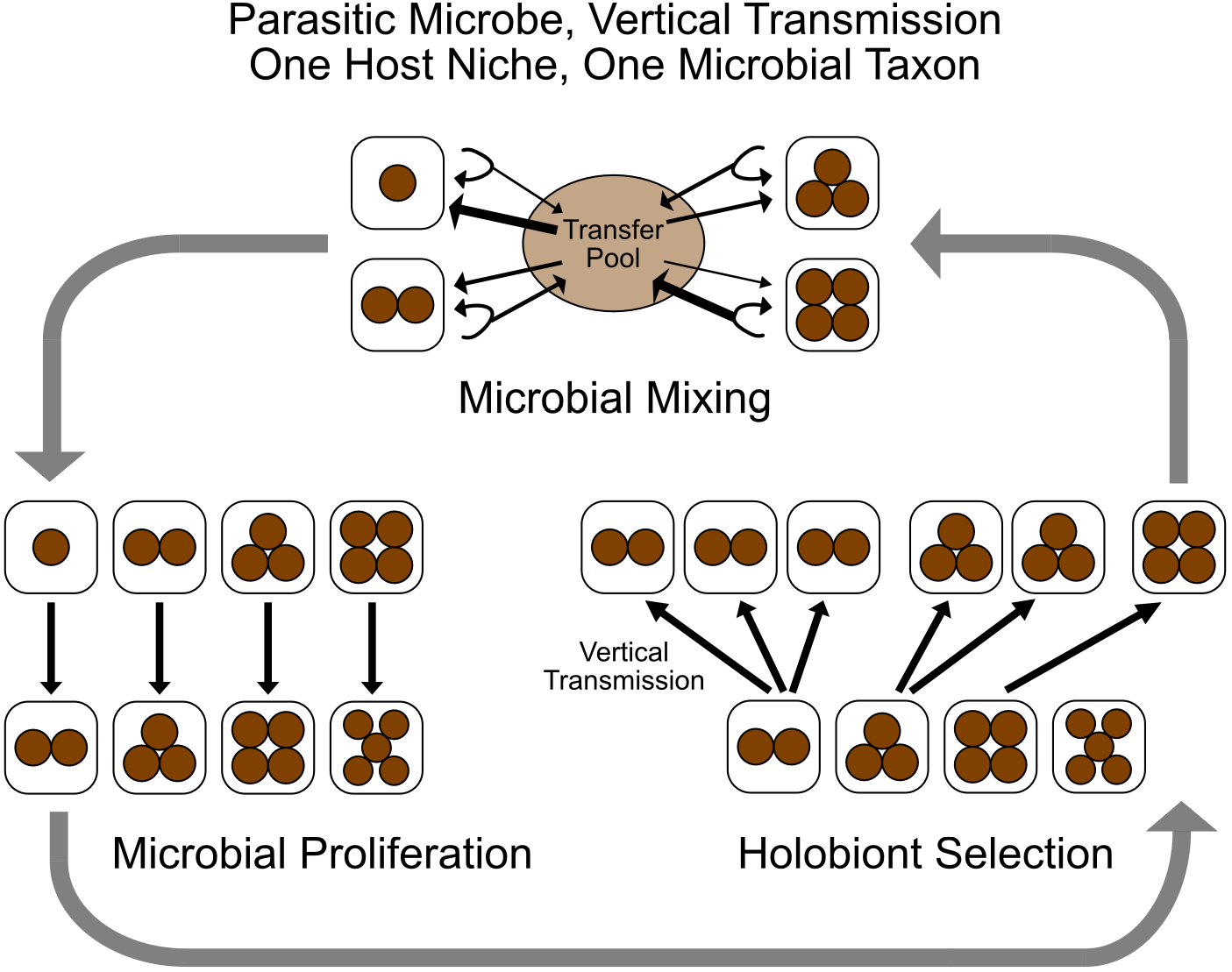
Sequence of processes with vertical transmission of parasitic microbiome.

**Figure 3:**
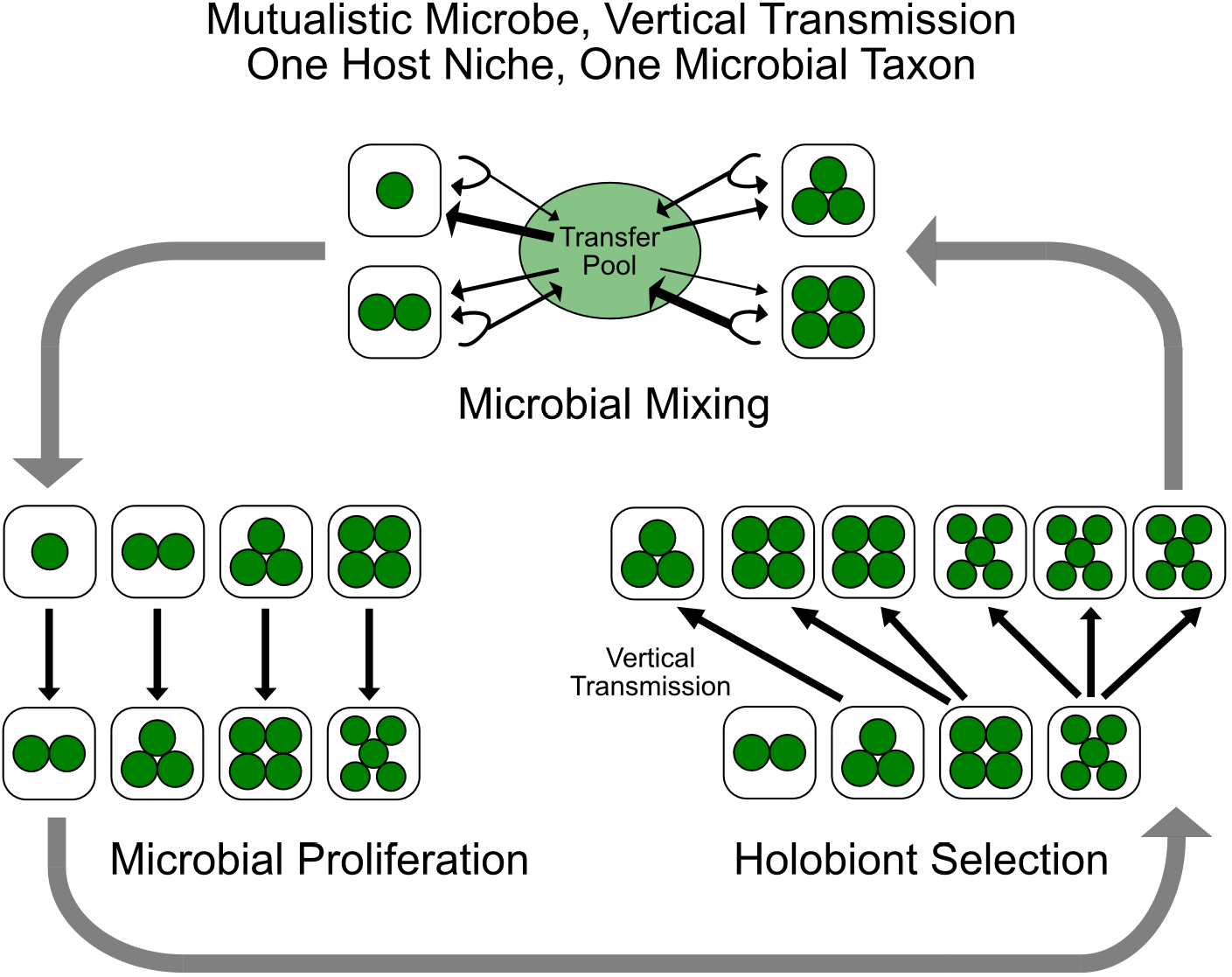
Sequence of processes with vertical transmission of mutualistic microbiome.

### Model

#### Microbial Mixing

Within a macro time step, the first stage is microbial mixing, as diagrammed at the top of Figures 2 and 3.

Each microbe has a small probability, *m*, of exiting its holobiont and entering a temporary “transfer pool.” Then the total transfer pool is divided equally across the holobionts. Each holobiont receives the same number of transfers. Hence, holobionts that had few microbes to begin with receive a net increase in microbe number and holobionts with many microbes to begin with incur a net decrease.

A small *m* is consistent with limited microbe interchange across hosts. The holobiont population needs limited horizontal mixing even though parents primarily transmit microbes vertically to their offspring. Otherwise, if *m* is 0, then holobiont selection reduces to clone selection on the hosts whereby the host clone with lowest number of parasites or the largest number of mutualists becomes fixed. Furthermore, if *m* is 0, then empty hosts cannot be colonized.

#### Microbial Proliferation

After microbial mixing, the next stage is for the microbes to proliferate within the holobionts, as diagrammed on the left side of Figures 2 and 3.

The microbes within each holobiont increase according to a density-dependent ecological population model. A single microbe strain obeys the logistic equation and multiple competing strains follow the Lotka-Volterra competition equations.

#### Holobiont Selection

Following microbial proliferation in the holobionts, the next stage is for the holobionts to be processed by holobiont selection, as diagrammed on the right side of Figures 2 and 3.

Each holobiont reproduces as a whole. The number of progeny a holobiont produces depends on the number of microbes in it, which is its holophenotype. Parental holobionts transmit their microbes to their juveniles such that the number of microbes in each juvenile equals the number of microbes present in its parent. In the case of microbial parasites in Figure 2, the holobionts with the lowest number of microbes leave the largest number of progeny and conversely for microbial mutualists in Figure 3. Holobiont reproduction and survival are independent of the density of holobionts.

The macro time step ends after the holobiont selection stage, whereupon the holobiont population is ready for another macro time step that begins once again at the mixing stage. Iteration of the macro time step with these three stages generates a trajectory of holobiont abundance and hologenotype frequencies through time.

The mathematical appendix includes instructions for how to download the model’s computer program from the website, GitHub. Also, the appendix includes a link to a short five-minute video on YouTube illustrating how to run the computer program.

### Results

Figures 4 and 5 show screen shots of the histogram through time of the hologenotype frequencies for host interactions with parasitic and mutualistic microbes. In both figures iteration of macro time steps leads from an initial uniform distribution to a stationary distribution of hologenotype frequencies at which holobiont selection balances microbial mixing.

**Figure 4:**
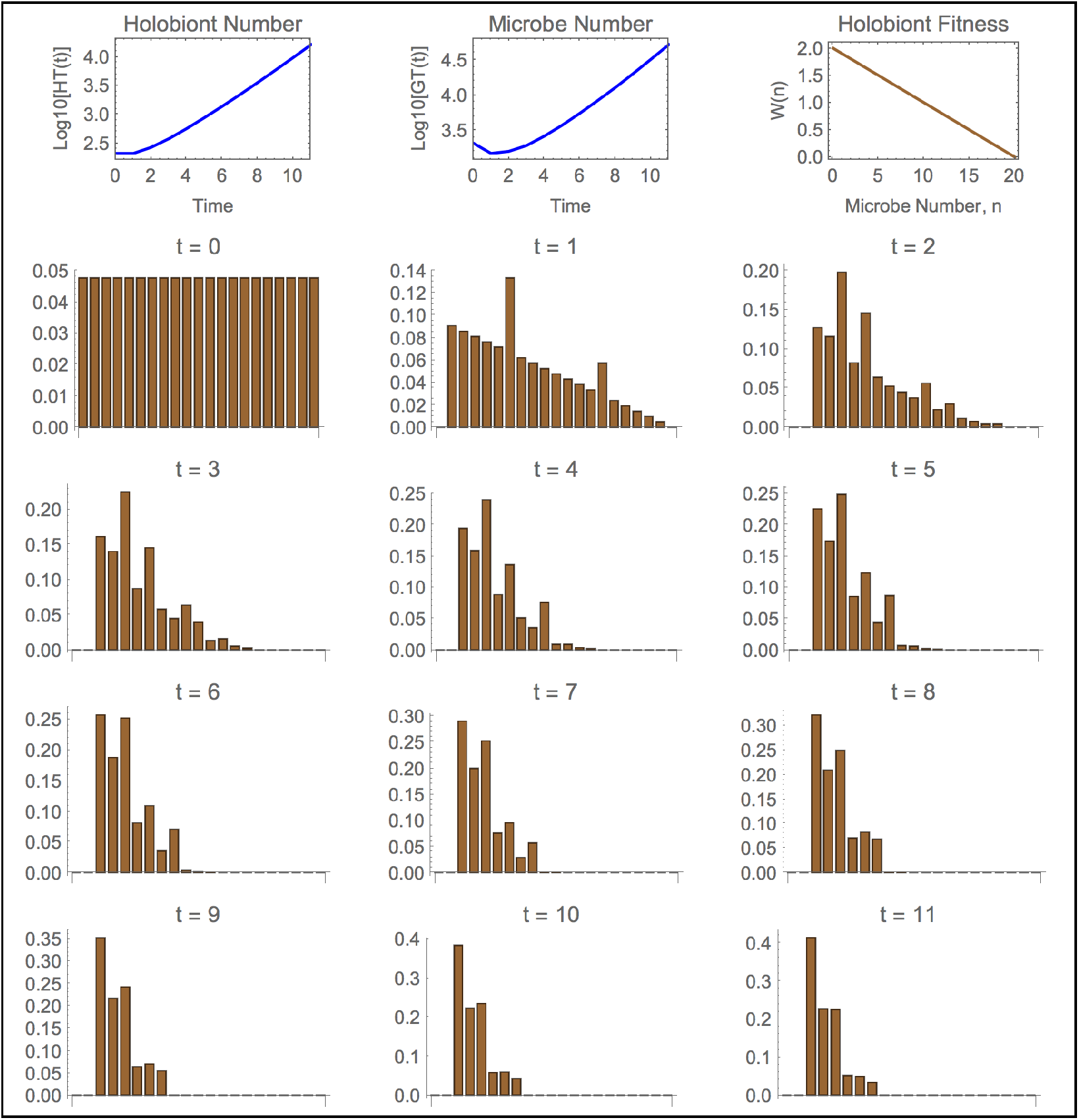
Results with vertical transmission of parasitic microbiome. Top left and middle graphs show log of total holobiont and microbe numbers through time and top right graph shows holobiont fitness as function of number of microbes in host. Histograms show distribution of hologenotype frequencies through time. The horizontal axis in the histograms is number of microbes per host and vertical axis is fraction of holobiont population.

**Figure 5:**
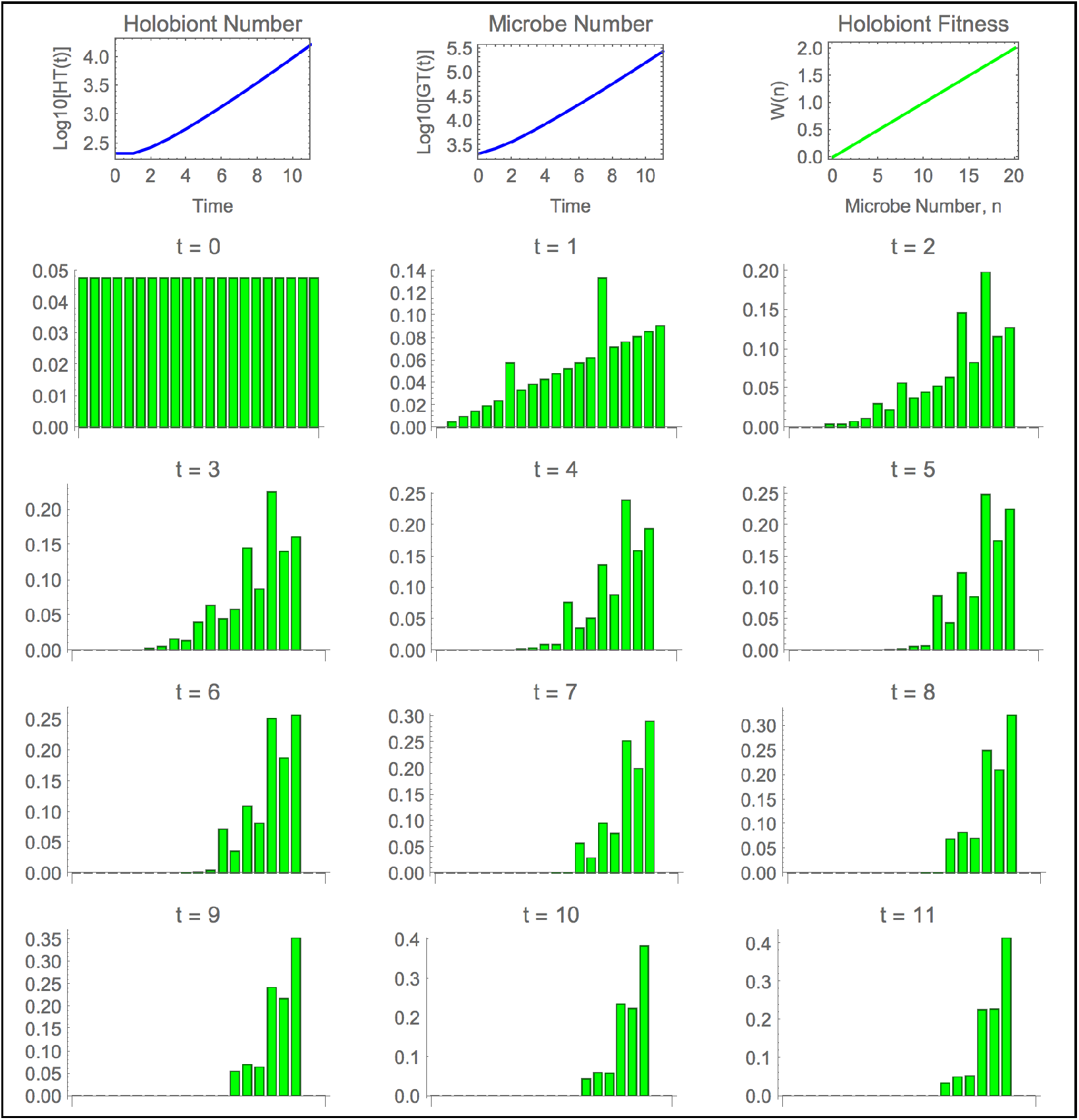
Results with vertical transmission of mutualistic microbiome.

In Figure 4 holobiont selection leads to a preponderance of holobionts with fewer parasites than initially. In Figure 5 holobiont selection leads to a preponderance of holobionts with more mutualists than initially. In the model’s picture display (not shown), the holobionts with a parasitic microbiome change through time from 50% brown to almost no color and holobionts with a mutualistic microbiome change through time from 50% green to nearly solid green.

The stationary distribution of hologenotype frequencies partly depends on the initial condition. An historical accident affecting the initial condition may influence the outcome of holobiont selection in this scheme of microbial mixing.

Microbial mixing homogenizes the hologenome and retards the progress of holobiont selec-tion, reinforcing an intuition that effective holobiont selection requires faithful vertical transmission of microbes with limited horizontal mixing.

For parasites, microbial proliferation acts in opposition to holobiont selection. By increasing the microbe’s intrinsic rate of increase and/or the number of micro time steps the virulence of the parasite is increased. This increased virulence can overtake holobiont selection, leading to holobiont extinction.

These findings support comparable conclusions about the evolution of gut microbiota via host epithelial selection (Schluter & Foster 2012).

In reality, much of the microbiome does not colonize vertically. Thus, another variant of the model is needed. One possible modification to the model is to allow the mixing probability, *m*, to equal 1, implying that every microbe leaves its host to enter the transfer pool only to be redistributed in equal measure back into now empty hosts. However, that would instantaneously stock all the hosts with identical microbiomes, denying holobiont selection any hologenotypic variation to operate on. Instead, sampling during microbe transmission can be introduced to the model. This sampling produces hologenotypic variation across holobionts and leads to the following formulation.

## Horizontal transmission

### Model

Three processes occur as sequential stages within each host generation, as shown in the top panels of Figures 6 and 7. In detail:

**Figure 6:**
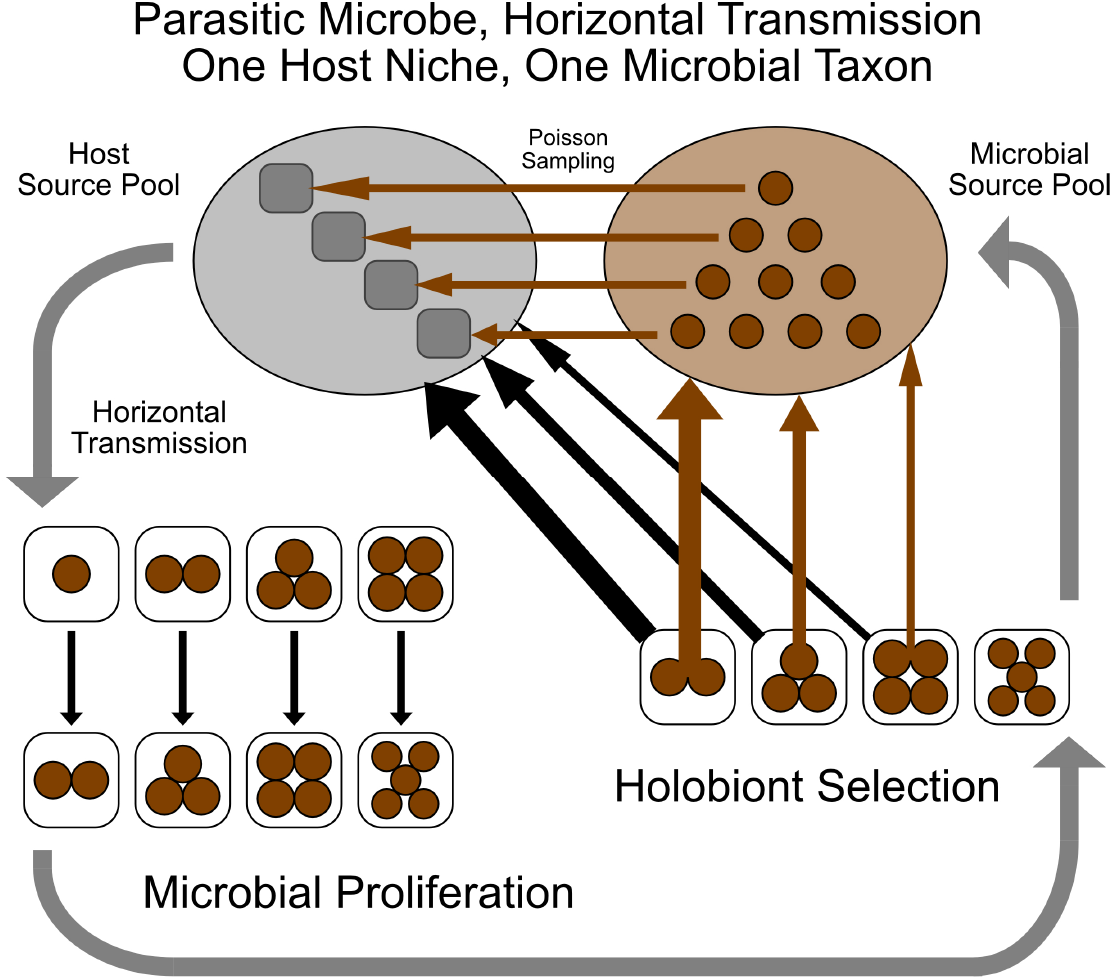
Sequence of processes with horizontal transmission of parasitic microbiome.

**Figure 7:**
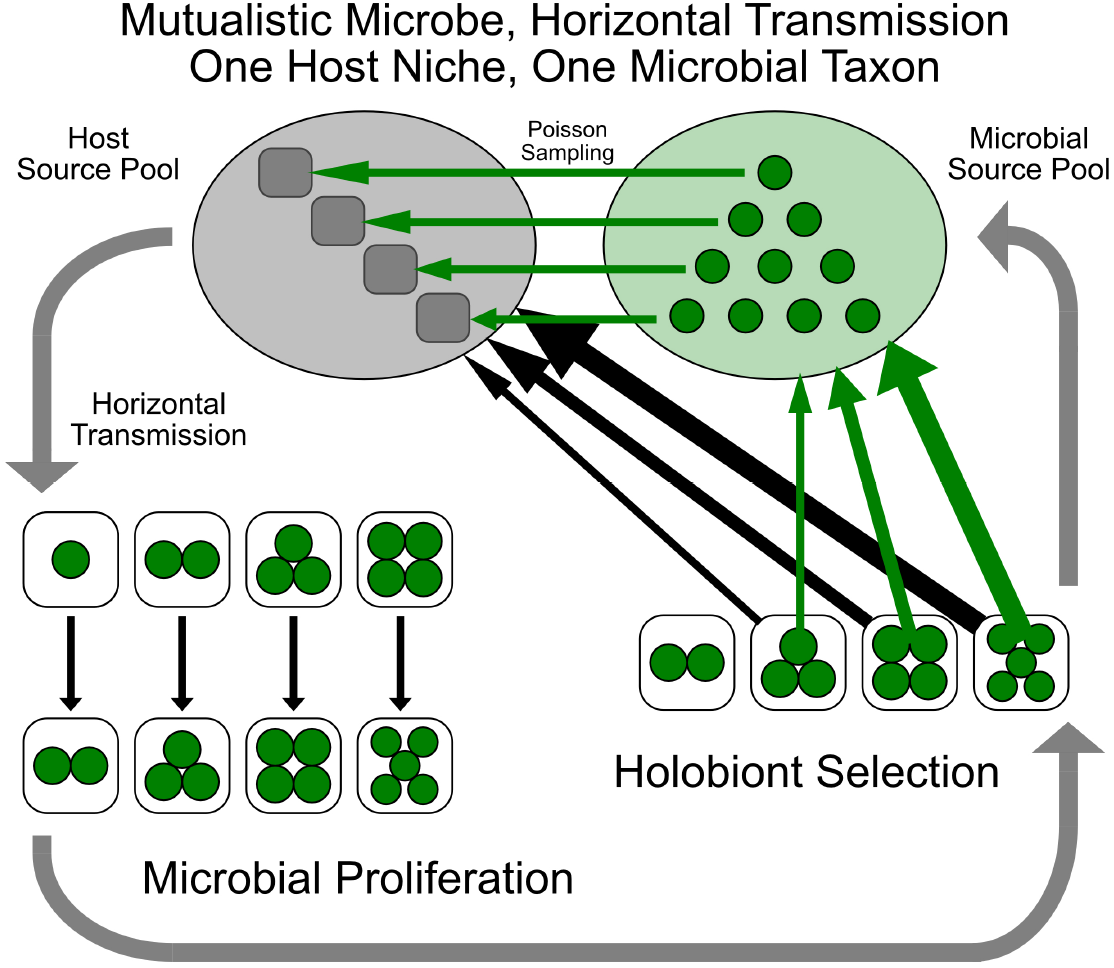
Sequence of processes with horizontal transmission of mutualistic microbiome.

#### Microbial transmission

Each generation begins with a normalized Poisson distribution of microbes across holobionts based on a density parameter equal to the number of microbes in its source pool divided by the number of juvenile hosts in its source pool.

#### Microbial Proliferation

The microbes within each holobiont increase according to density-dependent ecological population models, as before.

#### Holobiont Selection

Each holobiont reproduces as a whole. The number of progeny a holobiont produces depends on the number of microbes in it. Upon reproduction all the microbes produced from the holobionts enter a common microbe source pool and all the juvenile hosts join a common host source pool. For microbial parasites holobionts with the lowest number of microbes leave the largest number of progeny and conversely for the microbial mutualists. Holobiont reproduction and survival is independent of the density of holobionts.

### Results

Figures 8 and 9 show screen shots of the histogram through time of the hologenotype frequencies where the microbes are parasitic or mutualistic. Overall, the results are qualitatively similar in this model variant to those in the vertical transmission variant. Holobiont selection against parasites lowers average parasite microbe number in holobionts over generations relative to the initial condition and holobiont selection for mutualists increases average microbe number in holobionts over generations relative to the initial condition.

**Figure 8:**
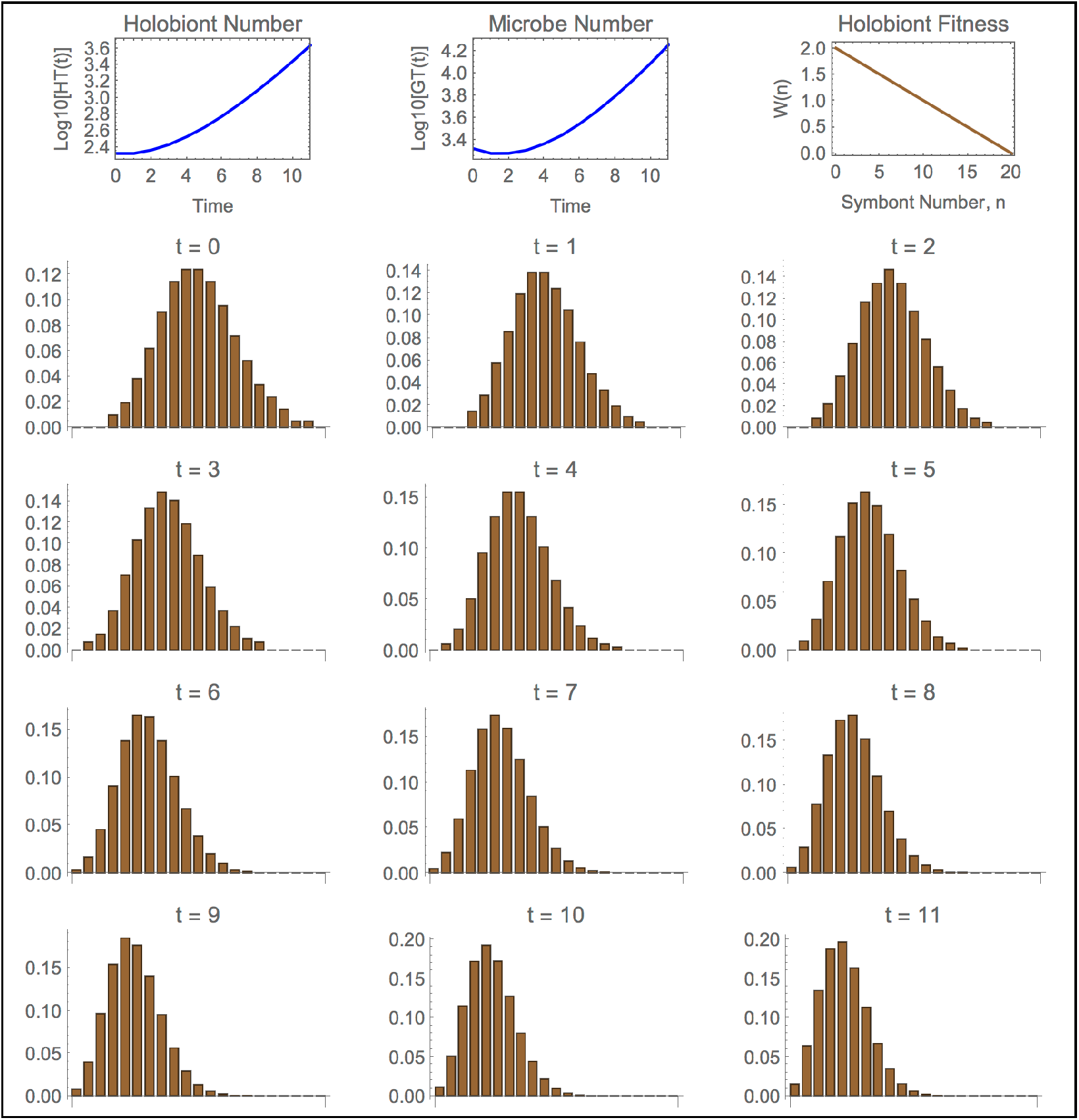
Results with horizontal transmission of parasitic microbiome.

**Figure 9:**
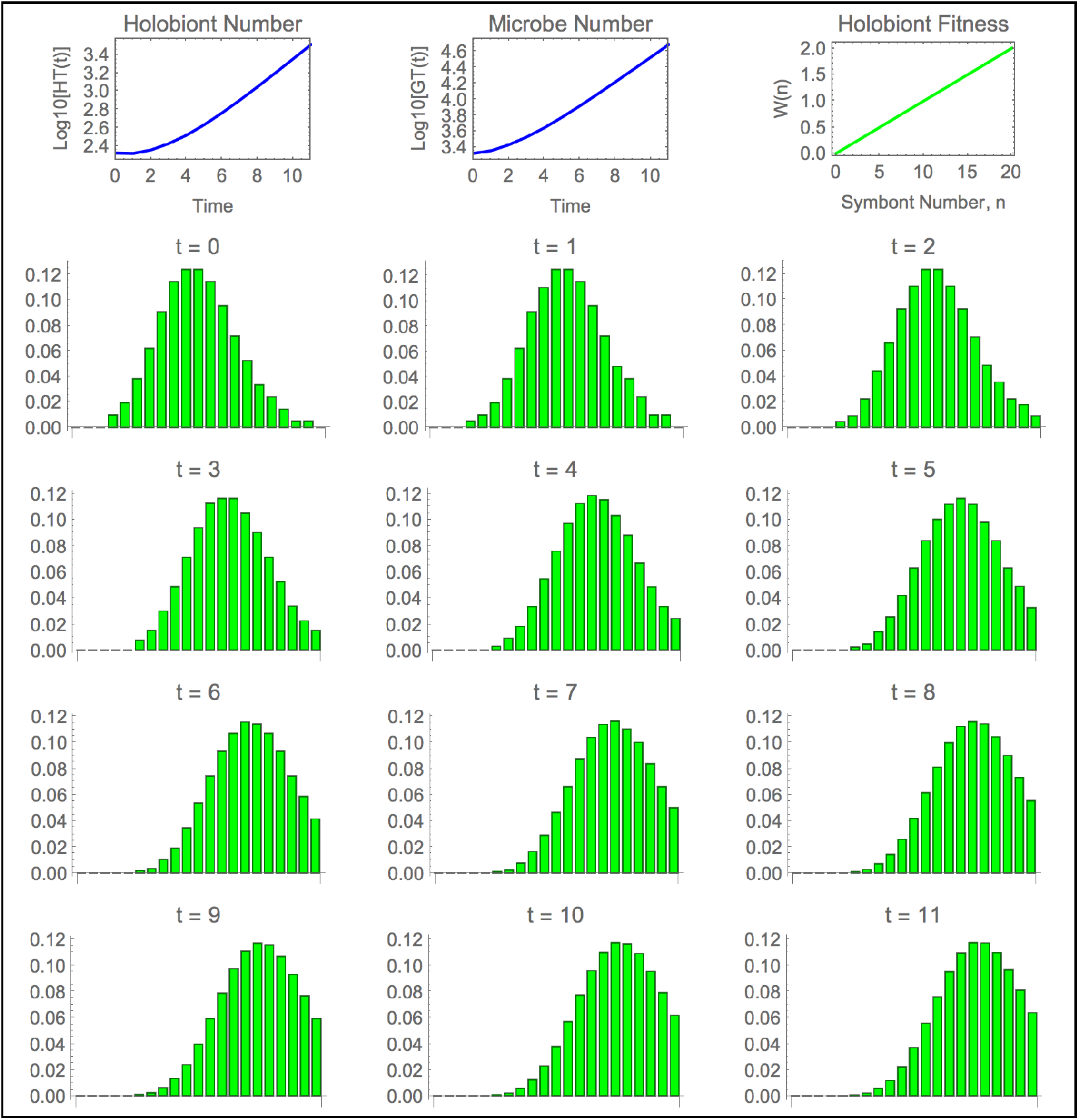
Results with horizontal transmission of mutualistic microbiome.

With horizontal transmission, a stationary distribution is attained when the holobiont selection balances the variability caused by the Poisson sampling of the source pools during transmission. (Whether this stationary distribution depends on initial conditions has not been analyzed.) Here too the virulence of the parasites can overcome the holobiont selection if the parasite intrinsic rate of increase is high enough and/or the number of micro time steps large enough.

## Discussion

The model here shows that holobiont selection causes the evolution of hosts with a reduced number of parasites and an increased number of mutualists, assuming either vertical or horizontal microbiome transmission. This outcome is to be expected given that reducing parasite load and increasing mutualists are adaptive. But *how* this expected outcome is attained is what makes holobiont selection distinctive. In fact, three types of evolutionary processes can be distinguished that can influence microbe abundance, one of which is holobiont selection.

First, parasite abundance may be lowered and mutualist abundance increased *via* host control. If the host has a locus in its nuclear genome which produces an antibiotic that kills parasitic microbes, then ordinary natural selection will fix alleles at that locus resulting in a reduced parasite load. Similarly, if the host has a locus that produces a chemical facilitating mutualistic microbes, then ordinary natural selection will fix alleles at that locus resulting in an increase of mutualists. A model in this framework has one equation—for changes in allele frequencies at a locus of interest in the host. This equation can be represented symbolically as a mapping from *p*(*t*) *→ p*(*t* + 1) where *p*(*t*) is an allele frequency at the locus of interest at time *t*. This evolutionary process does not acknowledge that the microbes are alive. In this process, parasitic microbes are no different than abiotic toxins and mutualistic microbes no different than abiotic nutrients. Therefore, host control does not seem to be a process capable of explaining the joint dynamics and evolution of the microbiome and host.

Second, the microbiome/host relationship can be influenced *via* coevolutionary selection. In a typical coevolutionary setup, the fitnesses influencing the gene frequencies in say, species-1 are functions of the abundances of both species-1 and species-2, and conversely for the fitnesses in species-2 (*cf.* Roughgarden 1983; Brown and Vincent 1987; Dieckmann and Law 1996; Carmona, Fitzpatrick and Johnson 2015). This setup leads to two coupled equations for changes in the gene pools of the two species which can be represented symbolically as a mapping from {*p*(*t*) *→ p*(*t* + 1), *q*(*t*) *→ q*(*t* + 1)} where *p*(*t*) and *q*(*t*) are allele frequencies in species-1 and species-2 at time *t*, respectively. The gene pools of the microbes and host are not merged and remain separate. The two species are playing an evolutionary game against each other, and the coevolutionary equilibrium is a Nash competitive equilibrium. This setup could be used to model how virulent a parasite should be taking into account the corresponding negative response by the host. Such coevolutionary selection would cause a net increase or decrease in microbe and/or host abundance depending on the feedbacks to their population dynamics. In a similar vein, mutualists who “cheat” by short-changing the benefits delivered to their host are thought to be repressed by host sanctions and other forms of retaliation (West, Kiers, Simms, and Denison 2002, Frederickson 2013). A mutualism may also be stabilized with reciprocal rewards (Kiers, Duhamel, Beesetty *et al*. 2011). In any case, the flavor of a coevolutionary account of mutualism is to emphasize a transactional “give and take” between antagonistic parties. This perspective has recently been extended to host/microbiome cooperation as well, where it is argued that “the host and each individual microbial strain are distinct entities with potentially divergent selective pressures” [p. 43] and further, that host control is needed to constrain the microbiome, to keep the microbiome on a “leash” (Foster, Schluter, Coyte and Rakoff-Nahoum 2017).

Third, the microbiome/host relationship can be influenced *via* holobiont selection, as has been demonstrated here. In holobiont selection, the gene pools of the microbes and host are merged into a single hologene pool and selection acts on a holobiont as a unit. The model consists of one “equation”, that is, the program to compute the vector of hologenotype numbers at time, *t* + 1, given the vector at time, *t*, is a single mapping that can be represented symbolically as *H*(*t*) *→ H*(*t* + 1). Thus, holobiont selection is a logical counterpart to ordinary natural selection, but based on hologenotypes rather than on traditional Mendelian genotypes. Moreover, in computer runs for cases such as those illustrated in Figures 4, 5, 8 and 9, the mean fitness of the holobionts monotonically increases through time, leveling off at equilibrium. One way to state the venerable fundamental theorem of natural selection is to say the mean fitness monotonically increases each generation until equilibrium, whereupon the mean fitness remains stationary (Karlin 1992, Frank and Slatkin 1992, Grodwohl 2018). The observation, according to computer runs of the model, that the mean holobiont fitness increases under holobiont selection until equilibrium is reached suggests that holobiont selection is similar to ordinary natural selection in this regard too. Like natural selection, holobiont selection is an adaptation-producing process. One might conjecture that a kind of fundamental theorem of holobiont selection will be found valid under similar assumptions to those that underlie the fundamental theorem of natural selection.

Whether holobiont selection is more important to holobiont evolution than coevolutionary selection is an open empirical question. The paper here shows that holobiont selection is a theoretically plausible evolutionary process. Holobiont proponents are therefore entitled to conjecture that holobiont selection is more effective in causing the evolution of microbiome/host integration than coevolutionary selection is.

The examples illustrate cases with a single microbe population, either a parasite or mutualist. If the microbiome includes two species, one a parasite and the other a mutualist, or two parasites of differing virulence or two mutualists providing different benefits, then the outcome is simply the combination of the outcomes from single species. Compare, for example, a nuclear genome with two unlinked non-epistatic loci. Selection simultaneously fixes the gene pool for the allele at each locus that confers the highest fitness. Similarly, if the parasitic and mutualistic microbes do not interact, then holobiont selection simply combines the outcome for both microbes, such as a simultaneous reduction in parasites and increase in mutualists. However, if the microbe species do interact with each other, then the Proliferation Stage in the model would need to incorporate population-dynamics for the species interaction instead of just the logistic equation presently used.

Although holobiont selection leads to the same qualitative outcome with both vertical and horizontal transmission, the underlying evolutionary processes are different between these two modes of transmission.

In vertical transmission, microbes incorporate into a host lineage along with the host’s nuclear genome. Microbial mixing loosens the association between microbes and the success of their holobiont’s lineage, thereby reducing the impact of the holobiont selection. Mixing also homogenizes the hologenotypes across holobionts erasing the hologenotypic variation that holobiont selection relies on to produce an evolutionary response.

In horizontal transmission, beneficial microbes from hosts that are successful because of having many mutualists that flood the microbial source pool. Hence, the next generation’s holo-bionts are assembled with an increase in microbe number because the ratio of microbes to hosts has increased. Conversely, the limited production of parasitic microbes from hosts that are successful because of having few parasites dilutes the microbial source pool. Hence, the next generation’s holobionts are assembled with a decrease in number of microbes because the ratio of microbes to hosts has decreased. The Poisson sampling of the microbes colonizing hosts guarantees hologenotypic variation for the holobiont selection to operate on.

With vertical transmission, a holobiont’s microbiome is inherited by *lineal descent*, albeit not as a Mendelian trait. In contrast, the process of microbiome assembly from the combined parental microbe production is dubbed here as *collective inheritance*.

Collective inheritance might prove valuable to understanding the evolution of bacteria and other microbes that feature large amounts of horizontal gene transfer (Mao & Lu 2016). It might also aid understanding cultural inheritance wherein offspring inherit a sample of the pooled knowledge of the parental generation.

A distinction between lineal and collective inheritance appears in the literature on multi-level selection in connection with the concepts of multi-level selection 1 (MLS1) and multi-level selection 2 (MLS2) (Mayo & Gilinsky 1987, Damuth & Heisler 1988, Okasha 2005, 2006). Okasha (2005) writes,

> “The key issue is whether the particles or the collectives constitute the ‘focal’ level, i.e., the level of interest. Are we interested in the frequency of different types of particle in the overall population of particles (which just so happens to be subdivided into collectives), or are we interested in the frequency of different types of collective themselves? If the former, then the collectives are significant only in that they constitute an environment for the particles, and thus may affect the particles’ fitnesses. We will judge evolution to have occurred when the overall frequency of different particle-types has changed. If the latter, then we are interested in the collectives in their own right, not simply as environments of the particles. We will judge evolution to have occurred when the frequency of different collective-types has changed. Damuth and Heisler (1988) refer to the first approach as multilevel selection 1 (MLS1), and the latter as multilevel selection 2 (MLS2).” [p. 1017]

Okasha (2006) further writes,

> “ In MLS2, collective fitness is defined as the number of offspring collectives. For this notion to apply, it is essential that the collectives reproduce in the ordinary sense, that is, they ‘make more’ collectives; otherwise, determining parent-offspring relations at the collective level will not exist. But in MLS1 this is inessential. The role of the collectives in MLS1 is to generate a population structure for the particles, which affects their fitnesses. For MLS1 to produce sustained evolutionary consequences, collectives must ‘reappear’ regularly down the generations, but there is no reason why the collectives themselves must stand in parent-offspring relations.” [p. 58]

Roughly speaking, holobiont selection with horizontal transmission corresponds with MLS1 and holobiont selection with vertical transmission corresponds with MLS2. But, holobiont evolution is not really the same as what these quotes are describing. In MLS1 the focus is on the microbes, with the host being secondary. Indeed, the host, or “collective” of MLS1 need not even be an organism. The collective need only be a kind of location, like the recurring haystacks in a field within which mice, the “particles”, can interact and realize their fitnesses in that context (Maynard Smith 1964). In contrast, with MSL2 the focus is on the host, with the microbes being secondary. The collectives give birth to more collectives depending on the particles in them while passing along their particles to their offspring collectives so both the collective and their particles share the same lineal parent-offspring relation.

Now, in holobiont evolution, both host and microbes are equally important assuming either mode of microbiome transmission, and neither is primary or secondary, because both the host and microbiome together comprise the holobiont itself and both participate in holobiont selection. Still, the distinction between MLS1 and MLS2 does foreshadow the distinction between lineal and collective inheritance encountered in the theory of holobiont evolution. Thus, although similarities between holobiont selection and multi-level selection exist, it may be fair to say that holobiont selection theory is not an extension of multi-level selection theory and that holobiont selection should be taken as a distinct process in its own right, *sui generis*.

Both lineal and collective modes of inheritance, although not named as such, are included in a recent model of microbiome evolution featuring both host and microbial selection (Zeng, Wu, Sukumaran & Rodrigo 2017). Further, the model here shares some similarity with literature on the evolution of organelles (Birky, Maruyama, & Fuerst 1983; Rand, Hane & Fry 2004; Smith 2007).

In a recent multilevel selection model for host/microbiome evolution, van Vliet and Doebeli (2019) concluded that “selection at the host level and in extension, at the holobiont level, could be of importance in nature but only under rather restrictive circumstances” [p. 5] including a requirement that “vertical transmission has to dominant over horizontal transmission” [p. 5]. The van Vliet-Doebeli model is an extension of a previous model described as a “mathematical formulation of Type II group selection” [p. 1569] (Simon, Fletcher and Doebeli 2012). As such, the van Vliet-Doebeli model is comparable to the vertical-transmission version of the model in this paper—its conclusions are not relevant to the horizontal-transmission version.

I have two remarks about the van Vliet-Doebeli paper. First, the result that holobiont selection is effective with a vertically transmitted microbiome only if the vertical transmission is not diluted too much by horizontal mixing has already been shown in (Roughgarden 2017, Roughgarden, Gilbert, Rosenberg et al. 2018) and is also mentioned in this paper when discussing the vertical-transmission case—but again, this concern does not pertain to microbial mixing in the horizontal-transmission case.

Second, van Vliet and Doebeli (2019) refer to a preprint of this paper, dismissing it with the claim that “by excluding variation in microbiome composition, this model could not address the question of under what conditions selection at the holobiont level can maintain traits that are disfavored by selection at the microbe level.” [p. 1]. This assertion is neither fair nor correct. After all, this paper is about vertical *vs* horizontal transmission—it did not intend to address the question of microbe/host tradeoffs. However, with this paper’s model, one *could* investigate a tradeoff between microbe and host fitnesses if one wanted to. Here’s how. Microbe fitness could be measured by its within-host carrying capacity, *k*, and the fitness of the host measured by, say, the slope of a line, *a*, relating holobiont fitness to microbe number. In the case of a parasitic microbe, one can ask for any pair, (*k*, *a*), if the microbe drives the holobiont to extinction. One can solve for curve in the *k × a* plane where coexistence of the parasite and host is just possible, such that points to one side of the line indicate coexistence and to the other side represent extinction. Additionally, for both a parasitic or mutualistic microbe, one can introduce a constraint equation relating the tradeoff in the microbe’s *k* to its effect on the host’s *a*. Then one solves numerically for the holobiont population growth rate, or equivalently, the holobiont mean fitness, once the stationary distribution of hologenotypes is attained. The particular tradeoff that corresponds with the highest holobiont mean fitness is the optimal tradeoff by the microbe to benefit the holobiont. The reciprocal problem could also be studied, whereby the host gives up some of its maximum fitness to increase the microbe carrying capacity. What is interesting mathematically in such an extension of this paper’s model is that the optimal traits for both microbe and host are found from the bivariate maximization of a scaler, namely, the holobiont mean fitness, whereas in coevolutionary selection, the optimal traits would involve vector maximization for separate microbe and host fitnesses using game theoretic methods. Actually, van Vliet and Doebeli’s (2019) assumption in their model that the host contains two types of competing microbes, “helpers” and “neutrals”, introduces unnecessary complications.

Previous investigators have noted that holobiont evolution is, in a sense, Lamarckian because holobiont selection affects acquired rather than inherited traits (Rosenberg, Sharon & Zilber-Rosenberg 2009, Osmanovic, Kessler, Rabin & Soen 2018). With horizontal transmission especially, a holobiont’s microbial abundance is acquired from environmental source pools. Still, evolution *via* holobiont selection seems best viewed as Darwinian rather than Lamarckian because holobiont selection is a selection process acting on holobionts as individuals. However, evolution *via* holobiont selection is not neoDarwinian because neoDarwinism is founded on Mendelian inheritance (Wright 1931).

The model here and its framework are extensible and scalable. Future theoretical research might investigate the comparative effectiveness of holobiont and coevolutionary selection in producing host/microbiome integration, how holobiont selection modulates microbiome community structure and how species diversity is maintained in the microbiome.

## Conclusion

The model here with its two variants demonstrates theoretically that holobiont selection is plausibly an important force in evolution because holobiont selection produces evolutionary change in holobionts with both vertical and horizontal microbe host transmission. Holobiont evolution can rely on collective inheritance as well as lineal inheritance.

## Acknowledgments

I am deeply grateful to Erol Akçay, Scott Gilbert, Priya Iyer, Ehud Lamm, Eugene Rosenberg, Jeremy Van Cleve, Vignesh Venkateswaran, and Ilana Zilber-Rosenberg for valuable comments on the manuscript. I thank two anonymous reviewers of this manuscript and an anonymous reviewer of a previous version for helpful comments.

## Appendix A: Mathematical Details

### State Variable Defined

The hologenotype numbers for a holobiont population through time are recorded in the matrix, *H*, where *H*(*t*, *n*) is the number of holobionts at time *t* containing *n* microbes. The top row of *H* records the initial condition of the holobiont population. The model computes the row at *t* + 1 given the row at *t*; that is, after each macro time step, the model appends a row to *H*. The index, *t*, runs from 1, which is the initial condition, to *t_M_* + 1 where *t_M_* is the number of macro time steps in the iteration. The index, *n*, runs from 0 to *K*, where *K* is the maximum number of microbes in a host. At the conclusion of *t_M_* macro time steps, *H* has become a matrix with dimensions, (*t_M_* + 1) *×* (*K* + 1).

The total number of holobionts through time is recorded in the vector,

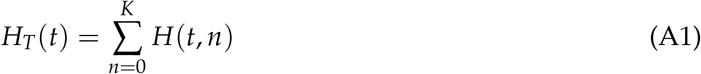

and the total number of microbes through time is recorded in the vector,

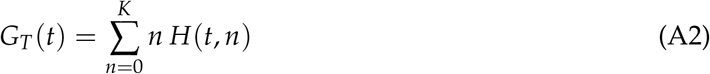

### Model Equations—Vertical transmission

Here are the three stages within a macro time step for the vertical-transmission variant of the model.

#### Horizontal Microbe Transfer Across Holobionts

The total number number of microbes that enter the transfer pool is *m G_T_*(*t*), where *m* is the probability a microbe leaves its host. Regardless of the host they reside in, all microbes have the same probability of entering the transfer pool. Therefore, the number of microbes who enter the transfer pool is simply *m* times the total number of microbes summed over all holobionts. The transfer pool is then distributed back equally across the holobiont population. The number of transfers received back from the pool per holobiont is then *m G_T_*(*t*)/*H_T_*(*t*), provided *H_T_*(*t*) *>* 0. Every holobiont receives this number.

Putting the arrivals and departures together, yields a formula for the new number of microbes, *n^t^* (*t*, *n*) in a holobiont that formerly contained *n* microbes at time, *t*.

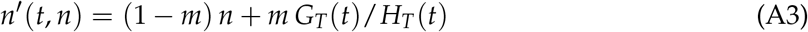

This formula says the new number of microbes in the holobiont equals the number there previously minus the number that departed plus the number that arrived.

The *n^t^* (*n*) must be an integer. To avoid considering fractional microbes, Equation (3) is rewritten as,

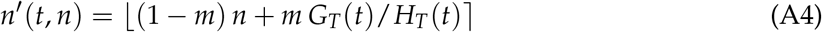

where the symbol ⎣*x*⎤ indicates the nearest integer to *x*.

The number of holobionts with *n* microbes after the horizontal transfer, *H^’^* (*t*, *n*), then is found by accumulating over *i* the holobionts whose microbe composition has changed from *i* to *n^’^*.

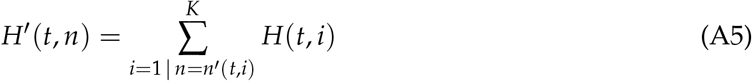

To see how this equation works, take a particular holobiont composition after horizontal transfer. Ask, for example, how many holobionts have exactly three microbes in them after the horizontal transfer has completed? *H^’^* (*t*, 3) is found by summing over *H* before the transfer started and using only the terms for the holobionts who wound up with 3 microbes after the transfer was over. If the summation is indexed with *i*, then the terms to include in the summation are those for which 3 = *n^’^* (*i*). These are the terms for holobionts who started with *i* microbes and wound up with 3 microbes. The same logic applies to *H^’^* (*t*, *n*) for any *n*.

#### Microbe Proliferation Within Holobionts

The next step is for the microbes to proliferate within their holobionts. If a holobiont starts with *n* microbes, then after one micro time step, the number of microbes in it is *F*(*n*), where *F*(*n*) is a demographic model for population growth.

The widely used logistic equation predicts sigmoid population growth with the population size leveling off at *K*, the carrying capacity. The speed of population growth initially, is *r*, the intrinsic rate of increase. Here, the model is written using a general *F*(*n*) to indicate that any suitable demographic model may be used, not necessarily the logistic equation.

Let *t_m_* be the number of micro time steps per macro time step. After *t_m_* steps, the microbe population will have grown in size to

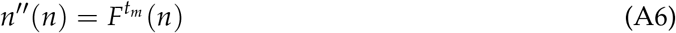

where 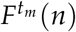 is the 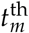 iterate of *F*(*n*). To avoid fractional microbes, the nearest integer is taken, yielding

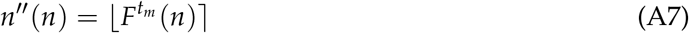

The number of holobionts with *n* microbes after microbe proliferation, *H^”^* (*n*), then is found using the same logic as in the preceding stage, by accumulating over *i* the holobionts whose microbe composition has changed from *i* to *n^”^*.

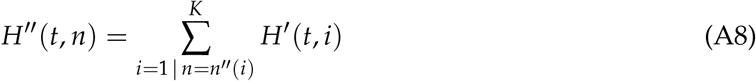

The microbe proliferation takes place in the holobiont population after the horizontal transfer stage. Hence, *H^’^* (*t*, *n*) appears in Equation (8).

#### Holobiont Selection

Here the number of microbes within a holobiont does not change, but the number of holobionts changes. The fitness of a holobiont with *n* microbes, defined as the number of progeny that the holobiont produces, is *W*(*n*). Therefore, the number of holobionts with *n* microbes after reproduction is simply

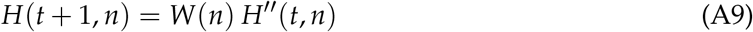

The holobiont selection takes place after microbe proliferation stage. Hence, *H^”^* (*t*, *n*) appears in Equation (9). Fractional holobionts are not meaningful, so the preceding equation is modified to indicate that the nearest integer is intended

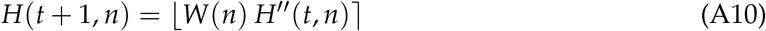

Overall, Equation (10) predicts the hologenotype numbers at time *t* + 1 based on what it was at time *t*.

### Model Equations—Horizontal transmission

Here are the three stages within a macro time step for the horizontal-transmission variant of the model.

#### Horizontal transmission From Source Pools

The macro time step begins with values for the size of the host pool, *H_T_*(*t*), and the microbe pool, *G_T_*(*t*). Then the Poisson density parameter indicating the average number of microbes per host is

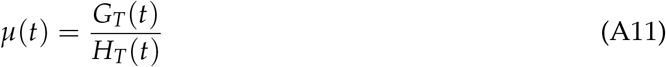

Next, the hologenotype numbers are found from a Poisson density function, *P_µ_*(*n*), normalized on the interval, [0, *K*], multiplied by the number of hosts at time *t*,

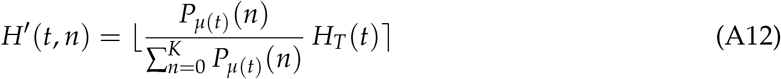

where 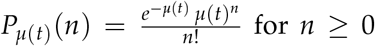. The numerator of the fraction in the equation above is the probability that an offspring of the host is colonized by *n* microbes given that the average number of microbes per host is *µ*(*t*) according to a Poisson probability distribution. However, the number of microbes in a host cannot exceed the host carrying capacity for microbes, *K*. So, the sum of probabilities of *n* from 0 to *K* is taken as the denominator in the fraction above. Thus, the overall probability of an offspring of the host being colonized by *n* microbes where *n* is between 0 and *K* is the fraction in the formula above. The sum of this fraction from 0 to *K* equals 1, as it should. This fraction then multiplies the number of hosts waiting to be colonized, *H_T_*(*t*), to yield the number of host offspring with *n* microbes at time *t*.

#### Microbe Proliferation Within Holobionts

As before, the next step is microbe proliferation within their holobionts. *F*(*n*) is a demographic model for population growth, such as the logistic equation.

After *t_m_* steps, the microbe population grows in size to

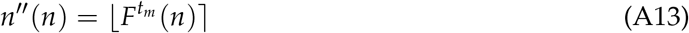

where 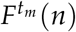 is the 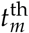 iterate of *F*(*n*).

The number of holobionts with *n* microbes after microbe proliferation, *H^”^* (*n*), is found by accumulating over *i* the holobionts whose microbe composition has changed from *i* to *n^”^*.

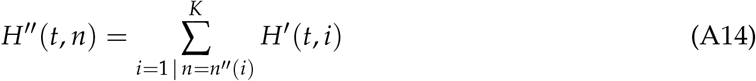

#### Holobiont Selection

The fitness of a holobiont with *n* microbes, defined as the number of progeny that the holobiont produces, is *W*(*n*). Therefore, the number of holobionts with *n* microbes after reproduction is simply

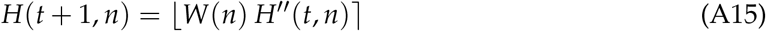

Prior to the next macro time step, all the microbes leave their holobionts and join a microbe source pool and the now-empty holobionts join a host source pool. Microbe retransmission and holobiont reassembly commences at the start of the next macro time step. The number of hosts and microbes in their source pools are, respectively,

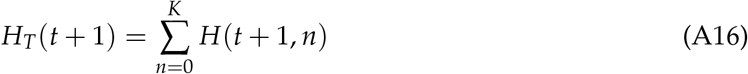

and

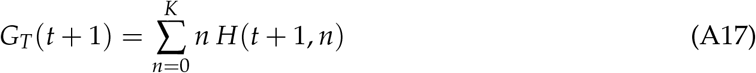

A new value of Poisson density parameter describing the average number of microbes per holobiont, *µ*(*t* + 1), is then computed for use at the beginning of the next macro time step.

### Download Program

Short programs written in *Mathematica* are available for download at: https://github.com/JoanKauai/Holobiont-Dynamics-Evolution.

The programs are supplied as both *Mathematica* Notebooks with an .nb extension and as Computable Document Format (CDF) files with an .cdf extension.

To use the .nb files one must have the *Mathematica* app which is quite expensive, although available in many universities. The .nb files allow the user not only to run the programs and also to adapt and change them by modifying the *Mathematica* code.

Using the CDF files requires a free CDF player that is downloaded from Wolfram at: https://www.wolfram.com/cdf-player/.

The player is analogous in principle to a PDF reader, only here CDF files rather than PDF files are displayed. The CDF files are fully interactive and one can change the model parameters with sliders. The only limitation to using CDF files rather than the notebook files is that one cannot modify the code of a CDF file.

If one is uncertain about whether to download the CDF player, one can use the CDF files directly online with a web browser. The vertical transmission version of the model is found at: https://www.wolframcloud.com/objects/joan.roughgarden/Published/HolobiontDynamicsVertical-05.cdf and the horizontal version at: https://www.wolframcloud.com/objects/joan.roughgarden/Published/HolobiontDynamicsHorizontal-05.cdf.

The CDF files run much faster when using the CDF player on a local computer, but clicking on these links provides a preview of how the programs work prior to deciding whether to download the player.

A short five-minute video demonstrating how the software works is available at https://youtu.be/19w7ZDkRe18

